# Actin-microtubule dynamic composite forms responsive active matter with memory

**DOI:** 10.1101/2022.01.10.475629

**Authors:** Ondřej Kučera, Jérémie Gaillard, Christophe Guérin, Manuel Théry, Laurent Blanchoin

## Abstract

Active cytoskeletal materials in vitro demonstrate self-organising properties similar to those observed in their counterparts in cells. However, the search to emulate phenomena observed in the living matter has fallen short of producing a cytoskeletal network that would be structurally stable yet possessing adaptive plasticity. Here, we address this challenge by combining cytoskeletal polymers in a composite, where self-assembling microtubules and actin filaments collectively self-organise due to the activity of microtubules-percolating molecular motors. We demonstrate that microtubules spatially organise actin filaments that in turn guide microtubules. The two networks align in an ordered fashion using this feedback loop. In this composite, actin filaments can act as structural memory and, depending on the concentration of the components, microtubules either write this memory or get guided by it. The system is sensitive to external stimuli suggesting possible autoregulatory behaviour in changing mechanochemical environment. We thus establish artificial active actin-microtubule composite as a system demonstrating architectural stability and plasticity.

---

Active materials are composed of a large team of energy-dissipating constituents. Local interactions of these constituents lead to an emergence of a collective self-organising behaviour ^1^, which has been widely studied with networks of cytoskeleton filaments in vitro ^2^. From the transient formation of patterns like asters and vortices ^3,4^ to constantly percolating active matter ^5–9^, which can be pre-programmed ^10^ or dynamically controlled ^11^, these synthetic out-of-equilibrium systems are considered to reconstitute, to a certain degree, the emergence of biological self-organisation ^2,12^. However, the ability of cells to develop, maintain, and adapt their internal organisation is not a mere consequence of the reorganisation of existing cellular components. Cells also dynamically assemble and disassemble their components. By this constant renewal of their cytoskeletal polymers, cells can maintain their architecture over longer periods, but they can also remodel it rapidly ^13,14^. Since the active cytoskeletal networks implemented up to now were made mostly of chemically stabilised polymers, they thus neglected fundamental dynamical properties that confer the living matter: its plasticity and adaptability.

In search of overcoming this major limitation, we wondered whether, in an in vitro assay, a combination of active self-organisation of cytoskeletal polymers with their dissipative self-assembly could constitute a more realistic life-like matter. Inspired by recent research on cytoskeletal cross-talk ^14,15^, we aimed at coalescing microtubules and actin filaments in a growing active composite. To do so, we developed a kinesin-driven motility assay of dynamic microtubules in the presence of growing actin filaments and a depletant. Unlike previously reported cytoskeletal composites, which relied on stabilised pre-assembled biopolymers ^16,17^, our system includes the self-assembly *in situ*, enriching the set of possible behaviours.

First, we tested the behaviour of each component separately. We attached kinesin-1 molecular motors to the passivated glass surface of the imaging chamber (Fig. 1 a) and flew in microtubule seeds that bind to the motors and glide in the presence of ATP. Free tubulin dimers added to the buffer together with GTP enabled the elongation of microtubules from the seeds. A crowding agent included in the buffer promoted cohesion of microtubules that are, otherwise, subject to electrostatic repulsion. In accordance with previous studies ^9,18^, the resulting attraction manifests by the transient formation of microtubule bundles (Fig. 1 b). These parallel and antiparallel bundles form as a result of the collisions of gliding microtubules (Fig. S1). The growth of microtubules translates into increased surface occupancy and increases the probability of their collisions. Consequently, the initially short, disordered microtubules evolve in an ordered network of long, quasi-aligned microtubule streams (Fig.1 b, c, Fig. S2, and Supplemental Movie 1), and so the global nematic order, a metric of orientation-ordering, grows until it saturates, preceding the saturation of microtubule assembly (Fig. 1 d and Fig. S3). Correlation analysis revealed that these streams form increasingly stable architecture (Fig. 1 e, f, Fig. S4) despite being supported by dynamic microtubules that stay motile within the streams as photobleaching experiments confirmed (Fig. 1 g, Supplemental Movie 2). Notably, the concentration of free tubulin and microtubule seeds in the assay has to exceed a certain threshold for the emergence of such stable orientation order (Fig. 1 h). The density of microtubules and thus the probability of their collisions is, therefore, central to the self-organising behaviour we observed.

**Fig. 1.**
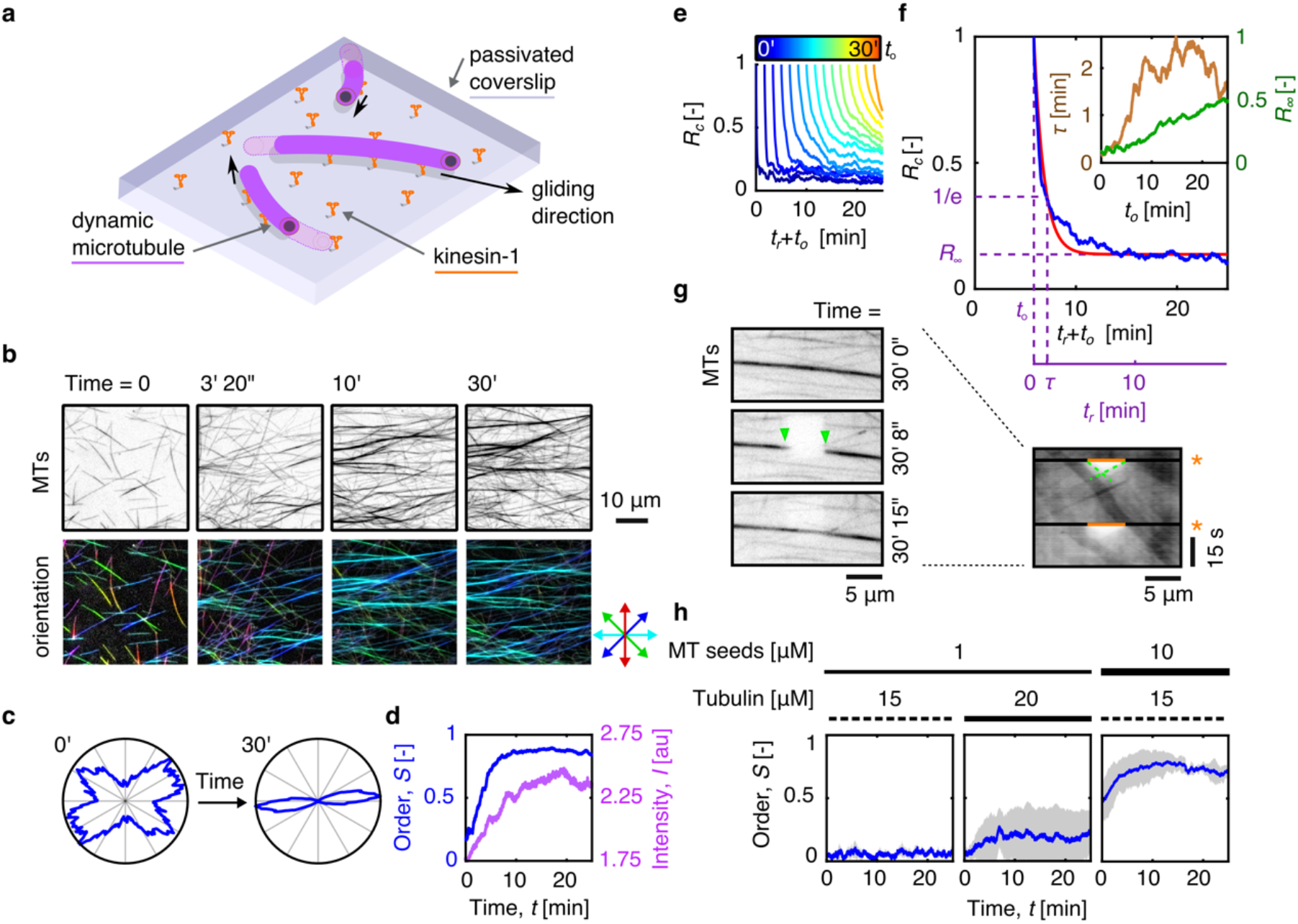
Motile dynamic microtubules system self-organises into quasi-parallel microtubule streams. **a**, Schematic representation of the system. **b**, Intensity inverted fluorescence micrographs of dynamic microtubule motility assay (top row) and the corresponding colour-coded orientation analysis (bottom row). **c**, Polar histogram of the orientation of microtubules is narrowing with time. **d**, Time trace of the global order, *S*, and fluorescence intensity, *I*, of dynamic microtubules in the motility assay. **e**, Time traces of equidistant vertical slices of Pearson correlation coefficient matrix of fluorescence micrographs of dynamic microtubule motility assay. **f**, The correlation slices in **e** can be approximated by an exponential decay with the parameters of the correlation decay time (time constant) and asymptotic correlation amplitude and represented by corresponding temporal profiles (inset). **g**, Intensity inverted fluorescence micrographs of photobleaching of microtubule stream (left, green arrowheads indicate the limits of photobleached area) formed during 30 minutes of ordering and the corresponding kymograph (right, orange lines indicate the photobleached area, green dashed lines indicate the front of receding photobleached areas, and asterisks denote the time of photobleaching). **h**, Time traces of the global order for various initial concentrations of microtubule seeds and free tubulin. All experiments were repeated at least four times with similar results. Data in **h** are represented as mean value (solid blue line) ± standard deviation (shaded area) (N = 4 per condition).

Next, we studied the ordering of growing actin filaments in the experimental chamber in the absence of microtubules (Fig. 2 a, Supplemental Movie 3). Actin filaments are not motile in our assay since they do not interact with kinesin motors. Crowding agent depletes actin filaments from the volume of the chamber to the glass surface, where a growing network forms locally nematic order (Fig. 2 b), similarly to previous report ^19^. As a result, in contrast to motile microtubules, the actin network alone does not show orientationally ordered architecture at a larger scale (Fig. 2 c, d and Fig. S5). Importantly, as the increasing surface occupancy reduces the mobility of the filaments, their organisation remains structurally stable, as the correlation analysis confirmed (Fig. 2 e). Varying the concentration of actin monomers within the range used in this study did not affect this emerging organisation significantly (Fig. 2 f).

**Fig. 2.**
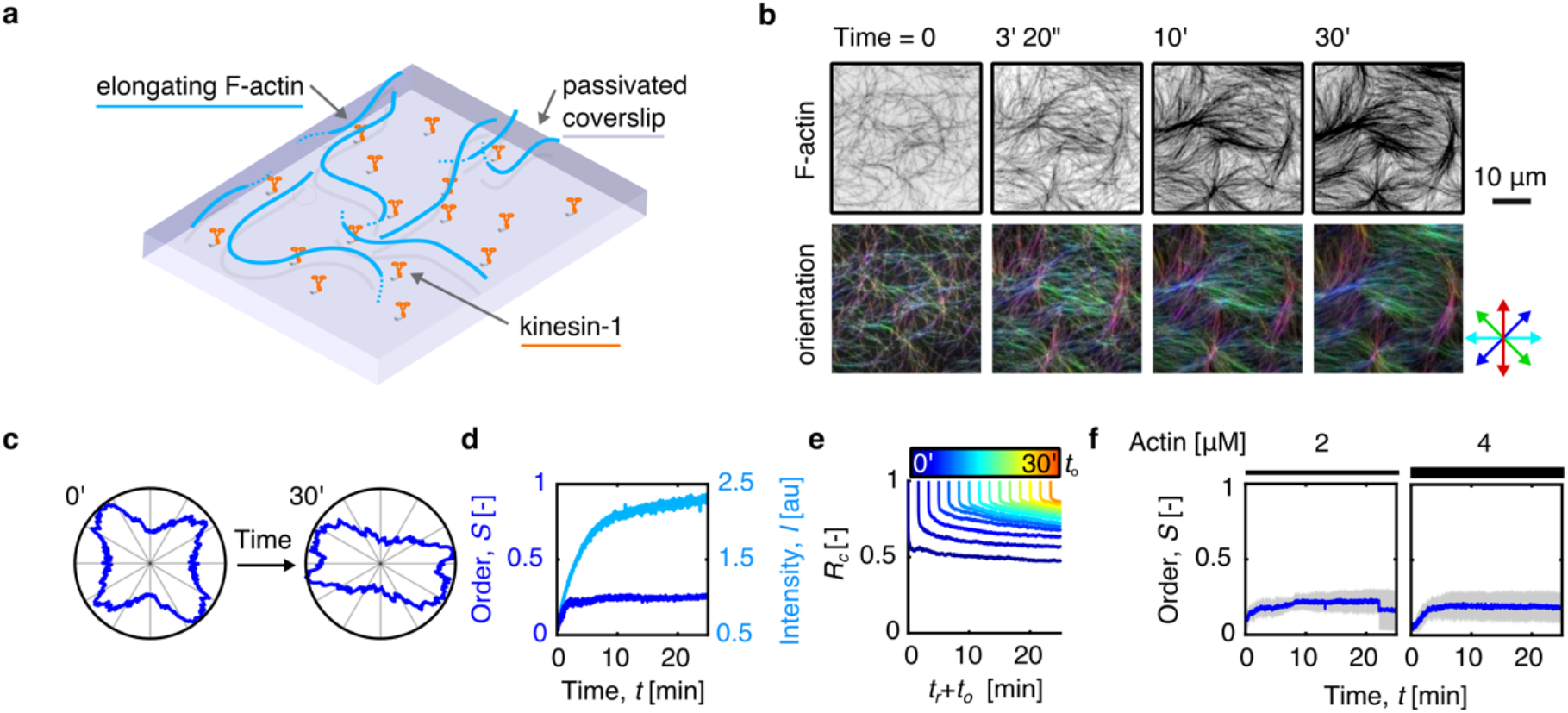
Assembling actin filaments self-organise in a locally nematic fashion with low global order. **a**, Schematic representation of the system. **b**, Intensity inverted fluorescence micrographs of dense elongating actin filaments (top row) and the corresponding colour-coded orientation analysis (bottom row) show local nematic ordering. **c**, Polar histogram of the orientation of actin filaments is narrowing only a little with time, indicating low global order. **d**, Time trace of the global order, *S*, and fluorescence intensity, *I*, of growing actin system. **e**, Time traces of equidistant vertical slices of Pearson correlation coefficient matrix of fluorescence micrographs of growing actin system. **f**, Time traces of the global order for two initial concentrations of free actin monomers. All experiments were repeated at least four times with similar results. Data in **f** are represented as mean value (solid blue line) ± standard deviation (shaded area) (N = 4 per condition).

After demonstrating that motile dynamic microtubules form self-renewing ordered architecture while growing actin forms a stable network, we merged these polymers in a composite system (Fig. 3 a) and studied how the interactions between them influence the ordering behaviour. Our central observation is that they do interact with each other. When microtubules encountered actin filaments, they occasionally caught the filaments and moved them temporarily, effectively organising the actin network (Fig 3 b and Supplemental Movie 4). More often, though, microtubules and actin filaments interacted cohesively. In such a case, denser regions of the actin network steered the microtubule trajectory, effectively guiding gliding microtubules (Fig 3 c and Supplemental Movie 5). These mutual interactions combine and constitute a feedback loop where microtubules organise actin filaments that in turn guide microtubules (Fig.3 d). Within a few minutes after commencing the experiment, this feedback in the dynamic composite system gave rise to an alignment between microtubules and actin filaments, which are forming overlapping streams and bundles, respectively (Fig. 3 e, Supplemental Movie 6). The reaching of this steady order preceded the saturation of the assembly of both cytoskeletal components (Fig. 3 f, Figs. S3-6). By comparing the structural stability of this composite with that of microtubules alone, we concluded that actin stabilises the emerging order (compare Figs. S3-6). An unexpected manifestation of this behaviour is that we could consistently observe the ordering of microtubules at a density that would not support ordering in the absence of an actin network (Fig. 3 g, Fig. S3).

**Fig. 3.**
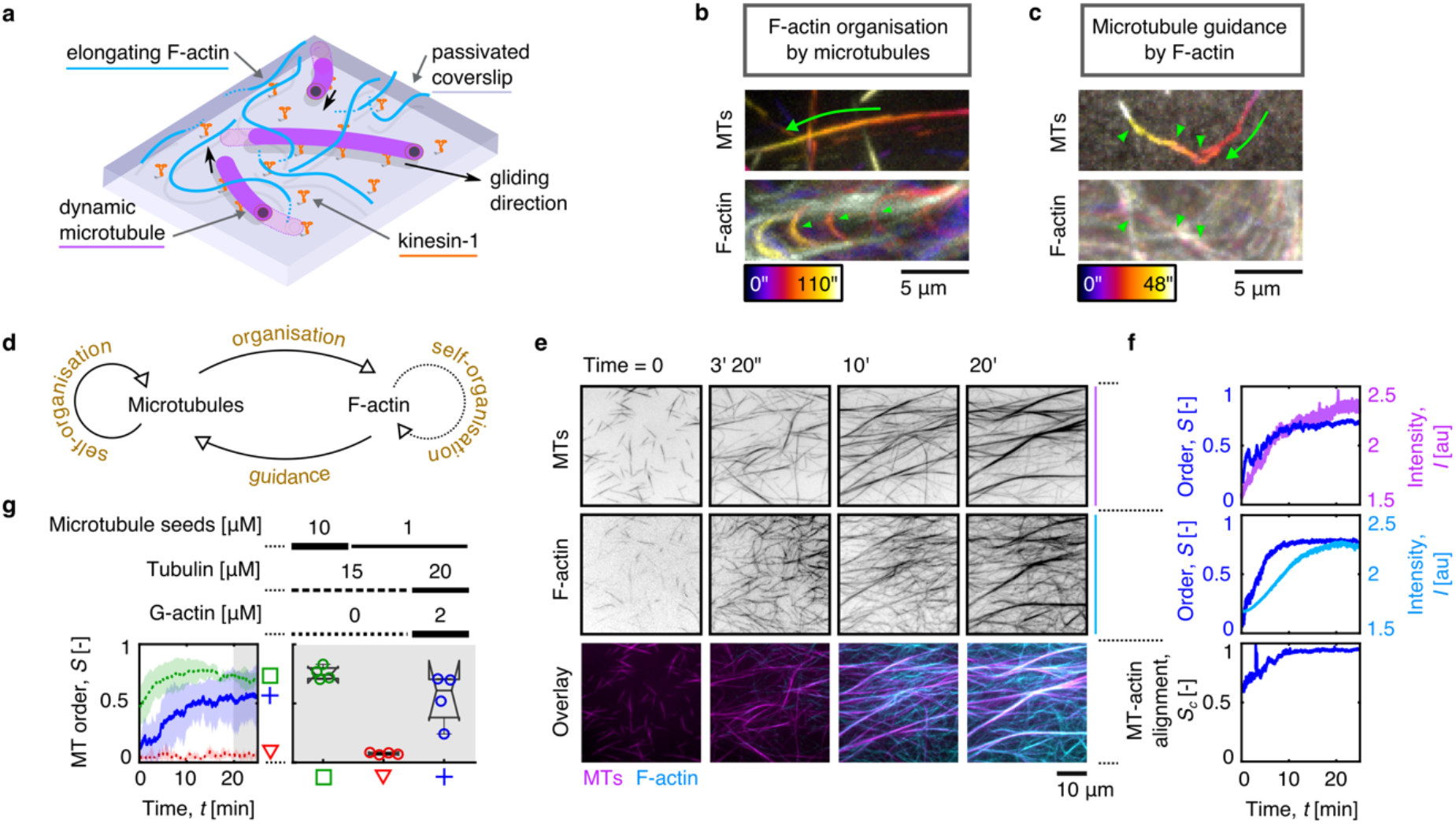
Interactions between microtubules and actin filaments mutually promote the self-organisation of the cytoskeletal composite. **a**, Schematic representation of the system. **b**, Colour-coded maximum fluorescence intensity projections of gliding microtubule (top) pushing actin filaments (bottom), effectively organising them. **c**, Gliding microtubule steered and guided by actin filaments. Image representation as in **b. d**, Schematic representation of the interactions forming a feedback loop between actin and microtubules. **e**, Multichannel fluorescence micrographs of self-assembling and self-organising actin-microtubule composite. Individual fluorescence channels are displayed as intensity inverted images. The overlay image is non-inverted for clarity. **f**, Time traces of the global order, *S*, and fluorescence intensity, *I*, of microtubules (top) and actin filaments (centre), and their mutual alignment, *S*_*c*_, (bottom) in the composite system. **g**, Time traces of the global order of microtubules (left) and the corresponding steady-state order (right) for three indicated initial biochemical conditions. All experiments were repeated at least four times with similar results. Data in **g** are represented as mean value ± standard deviation (left) and median ± 75^th^ percentile (right), where notches display the variability of the median between samples (N = 4 per condition).

To further investigate this unexpected emerging property of the composite, we tested how it would respond to the removal of one of its components. When we depolymerised microtubules either by adding CaCl_2_ or decreasing the temperature below 12°C, the actin network retained its ordering thanks to its cohesion and low turnover (Fig. 4 a, Supplemental Movie 7). Strikingly, when we, instead of microtubules, disassembled actin filaments using gelsolin, an actin severing protein, we observed microtubule dispersion (Fig. 4 b, Supplemental Movie 8). These observations highlight the role of the actin network in maintaining the order of microtubules over time.

**Fig. 4.**
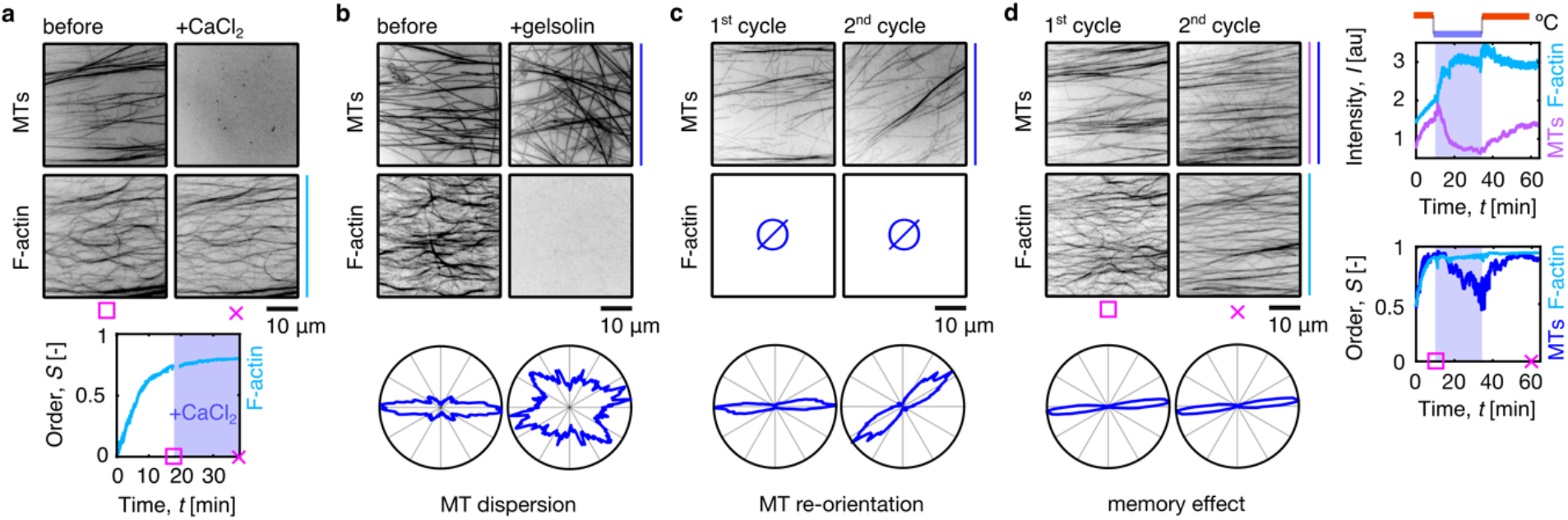
Response of the cytoskeletal composite to chemical and physical perturbations reveal the structural memory of the system. **a**, Intensity inverted fluorescence micrographs of microtubules (top) and actin filaments (centre) before (left) and after (right) the addition of CaCl_2_. Corresponding time trace of the global actin order (bottom). **b**, Inverted maximum fluorescence intensity projection (200 s) of microtubules (top) and intensity inverted fluorescence micrographs of actin filaments before (left) and after (right) the addition of gelsolin. Corresponding polar histograms of microtubule orientation (bottom) show microtubule dispersion. **c**, Intensity inverted fluorescence micrographs of microtubules (top) after the first (left) and second (right) cycle of microtubule polymerisation in the absence of actin. Corresponding polar histograms of microtubule orientation (bottom) show re-orientation in different directions. **d**, Intensity inverted fluorescence micrographs of microtubules (top) and actin filaments (centre row) after the first (left) and second (right) cycle of microtubule polymerisation. Corresponding polar histograms of microtubule orientation (bottom) show the retention of the orientation. Time traces of the fluorescence intensities (top right) and global order (bottom right) of microtubules and actin filaments. The shaded area corresponds to the cooling cycle. All experiments were repeated at least three times with similar results.

Importantly, these data suggest that actin acts as long-term structural memory for the microtubule network. We tested this hypothesis by reassembling microtubules after their depolymerisation. First, we let microtubules self-organise into streams in the absence of actin. After temperature-dependent depolymerisation, we re-polymerised the microtubules by increasing the temperature in the chamber. We observed that microtubules self-organised in streams again, but, importantly, the orientation of the streams was random and different from the initial orientation before the temperature shift (Fig. 4 c, Supplemental Movie 9). However, when we repeated this cycle in the presence of actin filaments, we witnessed a striking behaviour: the newly polymerised microtubule network recovered the ordering and orientation it had before disassembly (Fig. 4 d, Supplemental Movie 10). Therefore, microtubules could sense the order that was conserved in the organisation of the actin network. Actin filaments in our composite thus form a structural memory that acts as a template to sustain microtubule organisation ^20,21^.

The ability of the polymerised and ordered actin network to impose its order to microtubules reflects its structural stability. The stability stems from the low intrinsic actin turnover ^22^ and entropic cohesion due to the presence of a crowding agent ^23–25^, which practically limits the possibility of network remodelling. This is in sharp contrast with the initial phase of the composite self-assembly when the ordering and alignment of biopolymers occur. At this stage of lower surface polymer density, growing and gliding microtubules organise actin filaments into a steady conformation that will, in turn, guide microtubules. This feedback loop, together with the continuation of the polymerization, reinforces the spatial ordering of the composite, highlighting the importance of the self-assembling nature of the composite for its emerging properties. Notably, the order that is supported by the actin memory in our system is reversible as it can be annihilated by chemical signalling in the presence of regulatory proteins. Fine-tuning of the actin or microtubule turnover, which is intrinsic to biological systems, might thus constitute a means of precise tuning of the emergent order.

Structural memory of the cytoskeleton is a property used for cellular self-organization. Firstly, cytoskeletal filaments can support self-templated growth, for example, during filament guided filament assembly of the actin network in *C. Elegans* embryos ^21^. Secondly, one cytoskeletal network can serve as a template for the growth of another network, such as during microtubule guidance alongside actin bundles in growing neuronal tips ^26,27^, or along intermediate filaments ^20^. Although these two mechanisms likely combine, the latter can support a higher diversity of outcomes due to the combination of two populations of filaments with different lifetimes, specificity for regulatory factors, and mechanical properties. The system we presented here combines both types of structural templating observed in living matter but reserves the long-term memory for actin filaments, emphasising its composite character.

Introducing dissipative self-assembly of the constituting filaments to an active cytoskeletal composite is, therefore, critical for the emergence of adaptive architecture with concurrent plasticity and stability. While this feature is highly relevant for biological questions, it may also serve as a basis for developing diverse artificial systems, including life-like materials ^28,29^ and synthetic cells ^30,31^.

## Materials and Methods

### Protein production and purification

Actin and tubulin were purified and labelled as described previously ^32^. Briefly, bovine brain tubulin was isolated in temperature-dependent cycles of polymerisation and depolymerisation ^33^ and purified from associated proteins (MAPs) by cation exchange chromatography. Soluble tubulin was stored in the BRB80 buffer (80 mM Pipes pH 6.8, 1 mM EGTA, and 1 mM MgCl^2^) in liquid nitrogen. Rabbit skeletal muscle actin was purified from acetone powder ^34^. The actin was gel-filtered and stored in G-buffer (2 mM Tris-HCl pH 8.0, 0.2mM ATP, 0.5 mM DTT, 0.1 mM CaCl_2_, and 0.01% sodium azide) at 4°C. Parts of the tubulin and actin were labelled with Atto488 and Alexa568 fluorophores (Molecular Probes), respectively, by the NHS ester coupling ^35^ and stored as the unlabelled proteins. All experiments were carried out with 20% labelled tubulin and 5% labelled actin. Microtubule seeds were polymerised in the presence of GMPCPP. Recombinant, truncated kinesin-1-GFP motor and gelsolin were expressed in *E. coli* cells and purified similarly to previously reported methods ^36,37^ and stored at −80°C.

### In vitro assay

The in vitro gliding assay was performed in a flow chamber of approx. 15 uL, which was assembled from NaOH-cleaned glass coverslips using double-sided tape as a spacer. The channel’s surface was functionalised by GFP polyclonal antibodies (Invitrogen, A-11122) by filling the channel with 100 μg ml^−1^ antibodies in the HKEM buffer (10 mM HEPES pH 7.2, 5 mM MgCl_2_, 1 mM EGTA, and 50 mM KCl) for 3 minutes. The remaining available surface was then passivated by introducing 1% w/v bovine serum albumin (BSA) in HKEM buffer for 5 minutes. Next, kinesin-1-GFP motors (60 μg ml^−1^ in wash buffer: 10 mM HEPES pH 7.2, 16 mM PIPES buffer (pH 6.8), 50 mM KCl, 5 mM MgCl_2_, 1 mM EGTA, 20 mM dithiothreitol (DTT), 3 mg ml^−1^ glucose, 20 μg ml^−1^ catalase, 100 μg ml^−1^ glucose oxidase, and 0.3% w/v BSA) were specifically attached to the antibodies during 3 minutes of incubation. The channel was then perfused with wash buffer, and microtubule seeds (10 uM or 1 uM polymerised tubulin in wash buffer) were optionally introduced for 5 minutes of incubation, during which the seeds attached to kinesin motors. The unbound microtubules were washed away with wash buffer. Finally, the imaging Tic-Tac buffer (10 mM HEPES pH 7.2, 16 mM PIPES pH 6.8, 50 mM KCl, 5 mM MgCl_2_, 1 mM EGTA, 20 mM dithiothreitol (DTT), 3 mg ml^−1^ glucose, 20 μg ml^−1^ catalase, 100 μg ml^−1^ glucose oxidase, 2.67 mM ATP, 1 mM GTP, 0.3% w/v BSA, and 0.327% w/v methylcellulose (63 kDa, Sigma-Aldrich, M0387)) was introduced. Free tubulin (20 % labelled and 80 % unlabelled tubulin) and actin (12 % labelled and 88 % unlabelled actin) were added to the imaging Tic-Tac buffer optionally, with concentrations indicated in the main text. The flow chamber was optionally sealed by a capillary tube sealant (Vitrex) and transferred for imaging.

In the experiment with chemical actin and microtubule disassembly, the channel was kept open, and the environment was humidified to prevent evaporation. To induce actin disassembly, a 2 μl drop of 80 μM gelsolin and 10 μM tubulin mixture (to prevent microtubule depolymerisation) in the imaging Tic-Tac buffer was dripped to the channel opening. The proteins reached the imaging area by diffusion. To prevent flow-induced reorganisation of the cytoskeletal networks, no additional flow was used. After equilibration, the concentration of gelsolin was approximately 10 μM. To depolymerise microtubules chemically, ~ 10 μM final concentration CaCl_2_ was used. To depolymerise microtubules by a physical factor, the temperature was decreased to ∽12°C. Microtubules were re-polymerised by increasing the temperature to 37°C again.

### Imaging

Microtubules and actin filaments were imaged by total internal refraction fluorescence (TIRF) microscopy using an inverted microscope (Eclipse Ti, Nikon) with 100X 1.49 N.A. oil immersion objective (UApo N, Olympus). The Atto488 and Alexa568 fluorophores were excited by 491 nm and 561 nm lasers (Optical Insights), respectively, through the iLas^2^ dual laser illuminator (Roper Scientific). The fluorescence signals were separated by the Dual-View beam splitter (Optical Insights) and recorded by the Evolve 512 EMCCD camera (Photometrics). The sample was mounted to the environmental chamber maintained at 37°C and, optionally, at high relative humidity. To cool the sample down to depolymerise microtubules, the heating was turned off. The accumulated heat was transferred to a precooled bath, leading to a temperature of ∽12°C measured by a sensor attached to the microscope objective. The photobleaching of microtubule streams was performed using a UV laser at the end of the experiments. The imaging was controlled by the Metamorph software (v. 7.7.5, Universal Imaging), with images taken every second.

### Image processing and data analysis

Images were processed by FiJi (1.52v)^38^ and custom Matlab (v 9.6, MathWorks, Inc.) procedures using the MIJ ^39^ for running FiJi within Matlab. For presentation purposes, the contrast of images was adjusted. For analysis, the background of the 488-channel was subtracted using the built-in ImageJ function. The temporal profiles of the mean fluorescence intensities were obtained from ImageJ Plot Z-axis function. The kymographs were generated using the Multikymograph ImageJ plugin. As a measure of the orientation, we used the nematic ordering parameter, *S*, estimated from the mean resultant length of the director field ^40^. The value of *S* was calculated in Matlab from the weighted histogram of the orientation director field, which was obtained by OrientationJ plugin ^41,42^, as:

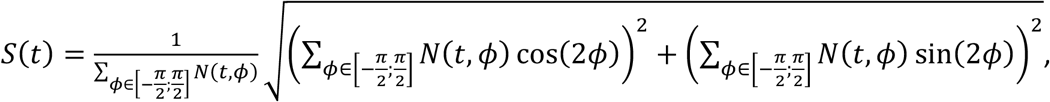

where *N* (*t, ϕ*) is the number of pixels oriented in the direction of angle *ϕ* at time *t* with coherency higher than 10 %.

The mutual ordering parameter was calculated from the orientation director, *O*_*MT*_ and *O*_*a*_, and the orientation energy, *E*_*MT*_ and *E*_*a*_, fields, which were obtained by the OrientationJ plugin, as:

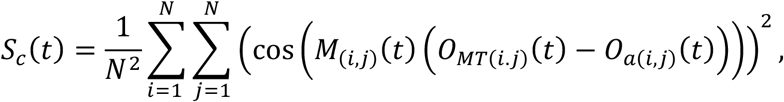

where (*i, j*) denotes the index within an image and the mask, *M*, is defined as *M*_(*i,j*)_(*t*) = 1 … *E*_*MT* (*i,j*)_ (*t*) > 0.05 ∧ *E*_*a* (*i,j*)_ (*t*) > 0.05 *M*_(*i,j*)_ (*t*) = 0 … otherwise.

To quantify the stability of the network organisation, the Pearson correlation coefficient of each frame of the experiment with all the consecutive frames was calculated. An exponential decay can approximate each series with two parameters: the correlation decay time (time constant), τ, and the asymptotic correlation amplitude. The correlation coefficients and curve fitting were calculated using built-in Matlab functions.

Graphs were produced using Matlab, and the final figures were formatted using Inkscape. At least three independent experiments were performed for each condition. No data were excluded from the study.

## Acknowledgements

This work was supported by the European Research Council (Consolidator Grant 771599 (ICEBERG) to MT and Advanced Grant 741773 (AAA) to LB. Our imaging platform is supported by the Laboratory of Excellence Grenoble Alliance for Integrated Structural & Cell Biology (LabEX GRAL) (ANR-10-LABX-49-01) and the University Grenoble Alpes graduate school (Ecoles Universitaires de Recherche, CBH-EUR-GS, ANR-17-EURE-0003).

## Author contribution

Conceptualization, OK, MT, LB; Methodology, OK, JG, CG; Investigation, OK; Formal Analysis, OK; Data curation, OK; Validation, OK; Resources, OK, JG, CG; Writing, OK, MT, LB; Visualization, OK; Supervision, MT, LB; Funding acquisition, MT, LB.

---

**Fig. S1.**
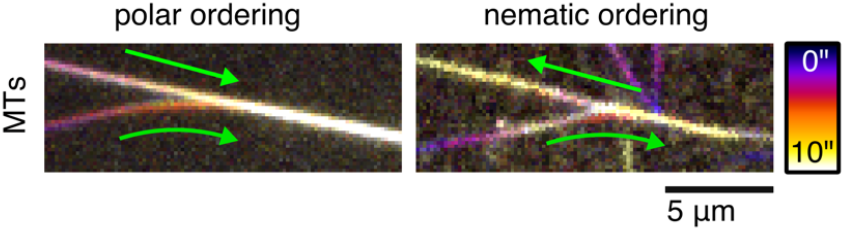
Colliding microtubules form a minimal stream. The collision of parallel microtubules leads to polar ordering (left). Collision of antiparallel microtubules leads to nematic ordering (left). Both types of streams can merge, forming a larger mixed stream (see Fig. 1 **g**).

**Fig. S2.**
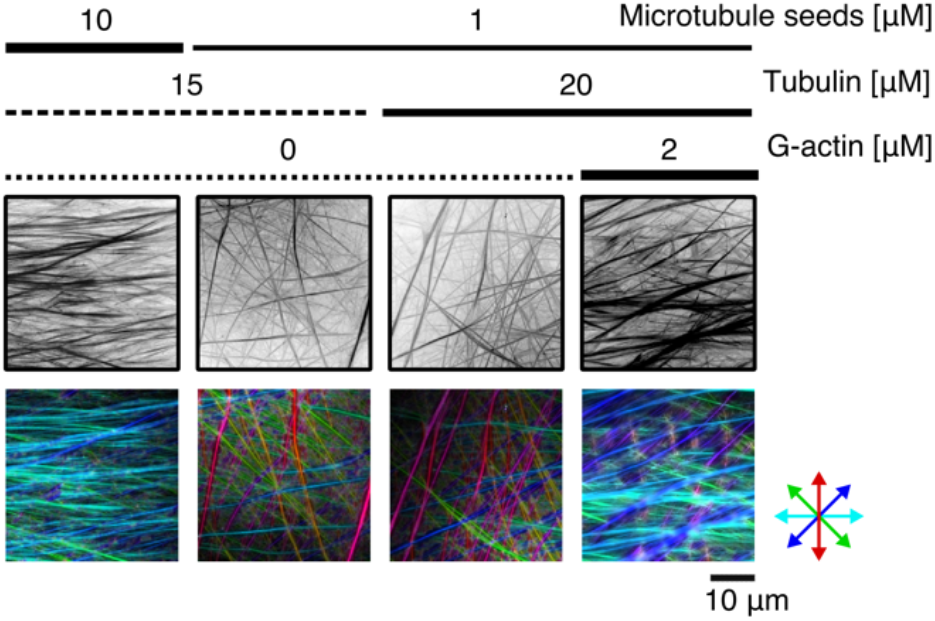
Microtubule ordering is concentration-dependent. Inverted maximum fluorescence intensity projection (30 min) of dynamic microtubule motility assay (top row) and corresponding non-inverted colour-coded orientation analysis (bottom row) for various initial concentrations of microtubule seeds, free tubulin, and actin monomers. All experiments were repeated at least four times with similar results.

**Fig. S3.**
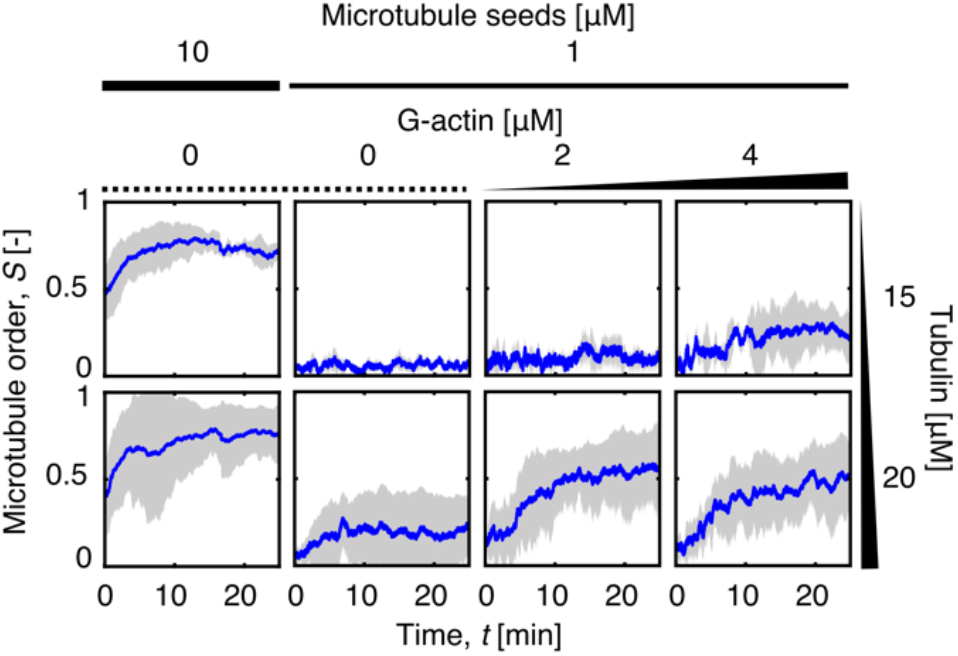
Microtubule global order is concentration-dependent. Time traces of the global microtubule order for all initial concentrations of microtubule seeds, free tubulin, and actin monomers used in this study. Data are represented as mean value (solid blue line) ± standard deviation (shaded area) (N = 4 per condition).

**Fig. S4.**
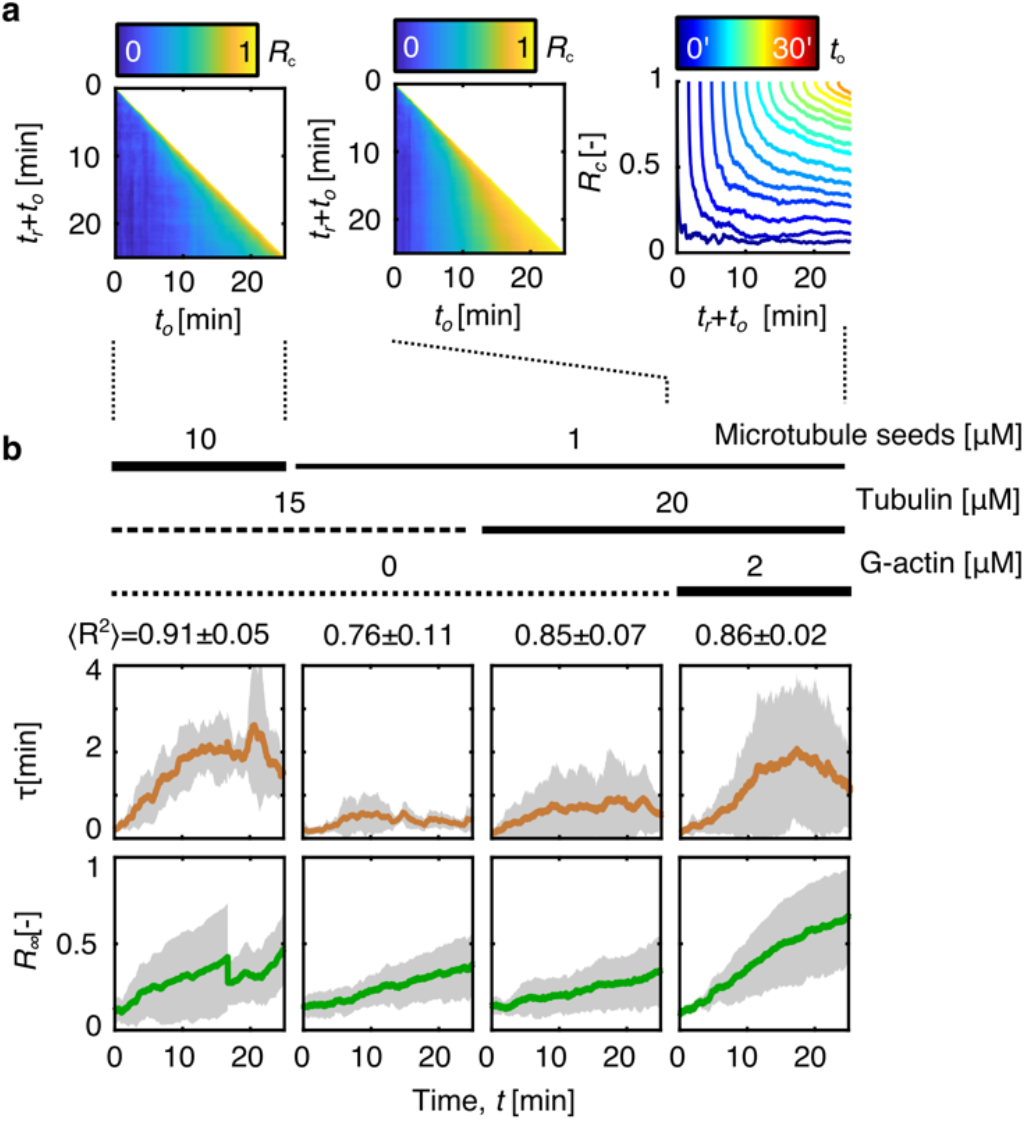
Correlation analysis. **a**, Examples of Pearson correlation coefficient matrix of fluorescence micrographs of dense, dynamic microtubule (10 μM tubulin in microtubule seeds, 15 μM free tubulin) motility assay (left) and composite system (1 μM tubulin in microtubule seeds, 20 μM free tubulin, 2 μM actin monomers, centre) with corresponding time traces of equidistant vertical slices of this matrix. **b**, Time traces of the correlation decay time (time constant) and asymptotic correlation amplitude of Pearson correlation coefficient matrix for various initial concentrations of the constituents. Data in **b** are represented as mean value (solid blue line) ± standard deviation (shaded area) (N = 4 per condition).

**Fig. S5.**
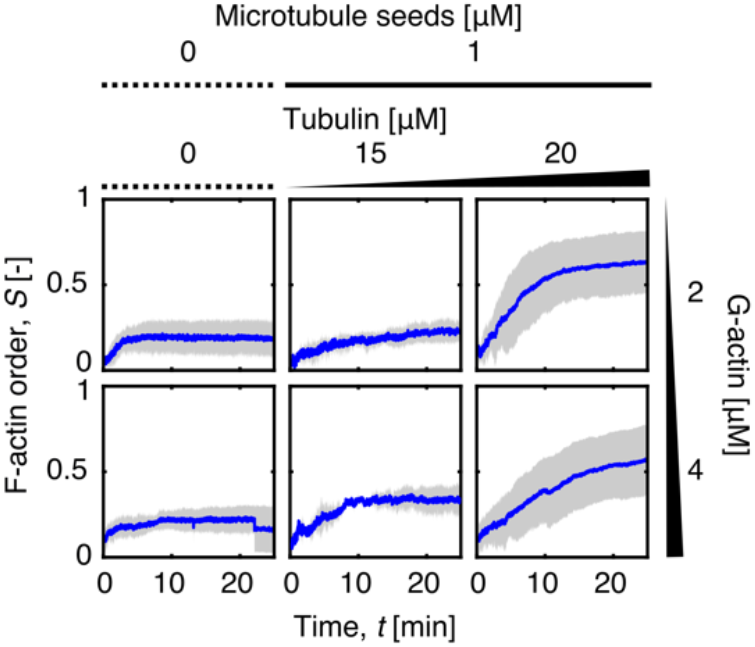
Actin global order depends on microtubule concentration. Time traces of the global F-actin order for all initial concentrations of actin monomers, microtubule seeds, and free tubulin used in this study. Data are represented as mean value (solid blue line) ± standard deviation (shaded area) (N = 4 per condition).

**Fig. S6.**
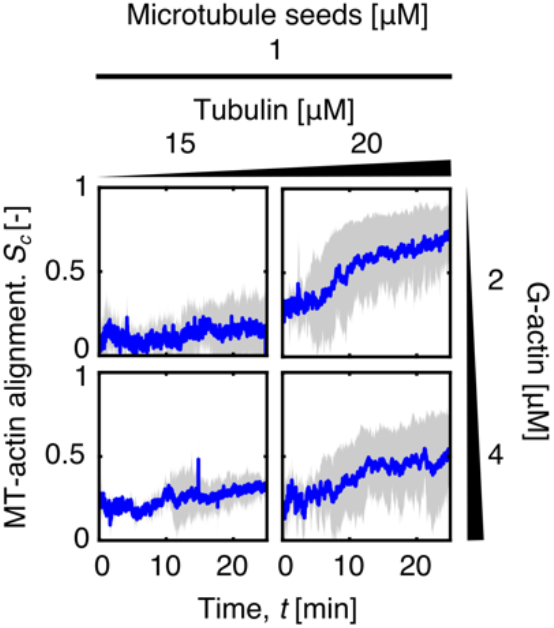
Microtubule-actin alignment. Time traces of the alignment of microtubules and F-actin for all initial concentrations of actin monomers, microtubule seeds, and free tubulin used in this study. Data are represented as mean value (solid blue line) ± standard deviation (shaded area) (N = 4 per condition).

## Supplemental Movies Captions

**Supplemental Movie 1. Motile dynamic microtubules system self-organises into quasi-parallel microtubule streams**. A sequence of intensity inverted fluorescence micrographs of dynamic microtubule motility assay (initial concentrations: 10 μM tubulin in microtubule seeds, 15 μM free tubulin).

**Supplemental Movie 2. Photobleaching of the microtubule stream reveals the conserved motility of microtubules within the stream**. A sequence of intensity inverted fluorescence micrographs of two consecutive executions of photobleaching of microtubule stream formed during 30 minutes of ordering.

**Supplemental Movie 3. Assembling actin filaments self-organise in a locally nematic fashion with low global order**. A sequence of intensity inverted fluorescence micrographs of dense elongating actin filaments (initial concentration: 2 μM actin monomers).

**Supplemental Movie 4. Gliding microtubule organises actin filaments**. A sequence of multichannel fluorescence micrographs of a gliding microtubule (magenta) pushing on actin filaments (cyan), effectively organising them.

**Supplemental Movie 5. Actin filaments guide gliding microtubule**. A sequence of multichannel fluorescence micrographs of actin filaments (cyan) deflecting a short gliding microtubule (magenta), effectively providing guidance.

**Supplemental Movie 6. Self-organisation of the cytoskeletal composite**. A sequence of multichannel fluorescence micrographs of a composite of growing actin filaments (cyan) and dynamic motile microtubules (magenta)(initial concentrations: 1 μM tubulin in microtubule seeds, 20 μM free tubulin, 2 μM actin monomers).

**Supplemental Movie 7. Retention of the order of actin filaments**. A sequence of multichannel fluorescence micrographs of microtubules’ (magenta) depolymerisation after the addition of CaCl_2_ in the presence of actin filaments (cyan).

**Supplemental Movie 8. Dispersion of ordered microtubules**. A sequence of multichannel fluorescence micrographs of microtubules’ (magenta) dispersion after the disassembly of actin filaments (cyan) by gelsolin.

**Supplemental Movie 9. Re-orientation of microtubules**. A sequence of intensity inverted fluorescence micrographs of dynamic motile microtubules undergoing a cycle of polymerisation, depolymerisation, and re-polymerisation in response to heating, cooling, and heating, respectively.

**Supplemental Movie 10. Memory effect in the cytoskeletal composite**. A sequence of multichannel fluorescence micrographs of a composite of growing actin filaments (cyan) and dynamic motile microtubules (magenta) undergoing a cycle of polymerisation, depolymerisation, and re-polymerisation in response to heating, cooling, and heating, respectively.

